# Single-cell profiling reveals a dose gap between population-average and complete pY-STAT1/3 suppression by JAK inhibitors

**DOI:** 10.64898/2026.10.26.759076

**Authors:** Karolina Ewa Zakrzewska, Jarosław Walczak, Karol Nienałtowski, Piotr Topolewski, Christina Maria Wagner, Agata Braniewska, Tomasz Rygiel, Richard Moriggl, Michał Komorowski

## Abstract

Strong suppression of a population-average signaling response need not place individual cells within the non-stimulated response range. We quantified this distinction in cytokine-induced JAK-STAT signaling using nuclear STAT1 phosphorylated at Tyr701 (pY-STAT1) and STAT3 phosphorylated at Tyr705 (pY-STAT3) in THP-1 and U937 cells exposed to six JAK-pathway inhibitors: solcitinib, BMS-911543, cerdulatinib, baricitinib, ruxolitinib, and tofacitinib. We compared arithmetic-mean inhibition and inhibition of the mean of log10-transformed single-cell responses (log-mean) with the fractions of cells leaving the stimulated response range or entering the non-stimulated range. Arithmetic-mean inhibition was left-shifted relative to log-mean inhibition, which more closely tracked departure from the stimulated range. Neither population summary uniquely specified entry into the non-stimulated range. At the lowest tested doses where fitted arithmetic-mean or log-mean inhibition reached 80%, median fractions remaining above that range were 76% and 37%, respectively. The apparent IC50 for complete-inhibition occupancy was a median 2.4-fold higher than the log-mean IC50 and 7.4-fold higher than the arithmetic-mean IC50. The size of this dose gap varied with cell line, inhibitor, and phospho-target within the tested panel. These results provide an empirical basis for interpreting conventional inhibition curves in terms of single-cell response fractions and show why population attenuation and complete-inhibition occupancy should be reported as distinct pharmacological endpoints.

## Introduction

Signaling pathways translate extracellular cues into cellular responses, making their pharmacological modulation central to the treatment of diverse diseases [1, 2]. Yet even apparently uniform populations can respond heterogeneously to the same stimulus. Such differences can reflect variation in molecular content and cellular state rather than simply noisy signaling [3–5]. In this setting, a decrease in the average response need not mean that most cells have returned to their non-stimulated response range.

Single-cell cytokine responses span broad, overlapping distributions [5, 6]. Population averages can therefore conceal biologically relevant fractions of responding cells, particularly when a minority retains elevated activity despite strong average suppression [4, 7]. Previous studies have shown that drug-response metrics can obscure minority states and qualitatively different modes of cellular inhibition [8–10]. What remains less clear is how conventional inhibition curves relate quantitatively to these cell fractions, and how this relationship varies across biological and pharmacological contexts.

This raises two distinct pharmacological questions: how strongly is the typical response attenuated, and what fraction of cells enters the non-stimulated response range? We use state occupancy to mean the fraction of cells within a predefined response range. Complete-inhibition occupancy denotes entry into the non-stimulated range, an operational criterion rather than proof that all downstream signaling has ceased.

We addressed these questions in cytokine-induced JAK-STAT signaling, using nuclear STAT1 phosphorylated at Tyr701 (pY-STAT1) and STAT3 phosphorylated at Tyr705 (pY-STAT3) as readouts. Interferon-*γ* (IFN-*γ*) predominantly activates STAT1, whereas interleukin-10 (IL-10) predominantly activates STAT3, through partially shared upstream JAK components [11–14]. These readouts allow comparison of related signaling routes without assuming that phosphorylation directly measures downstream function. We refer to inhibitor-treated cells remaining above the non-stimulated pY-STAT reference range as residual pY-STAT-positive cells.

The six-inhibitor panel provides pharmacologically diverse JAK-pathway perturbations. Solcitinib, BMS-911543, and the dual SYK/JAK inhibitor cerdulatinib have different reported target profiles; baricitinib, ruxolitinib, and tofacitinib extend the panel to clinically used JAK inhibitors with partly overlapping activity [15–17]. The purpose is to test whether the relationship between population suppression and state occupancy recurs across distinct perturbations, not to infer biochemical selectivity from pY-STAT responses.

In THP-1 and U937 cells stimulated with IFN-*γ* and IL-10, we compared inhibition of the arithmetic mean and of the mean of log10-transformed single-cell responses, hereafter log-mean, with two state-occupancy endpoints. Partial inhibition measures departure from the stimulated response range, including cells that have already entered the complete-inhibition range. This design tests how informative a population-inhibition curve remains when the endpoint is defined by a fraction of cells rather than an average response.

Here we establish an empirical relationship between conventional inhibition curves and the fractions of cells leaving the stimulated pY-STAT1/3 range or entering the non-stimulated range. Across six JAK inhibitors, two cell lines, and two readouts, the apparent IC_50_ for complete-inhibition occupancy was a median 2.4-fold higher than the log-mean IC_50_ and 7.4-fold higher than the arithmetic-mean IC_50_. These comparisons quantify both the dose gap and its variation across conditions, providing a basis for estimating how much single-cell responsiveness may remain when a population-inhibition curve indicates strong suppression.

## Results

### Single-cell response distributions motivate state-based inhibition metrics

To motivate how inhibitor effects should be quantified in this system, we first examined representative single-cell distributions of pY-STAT1 and pY-STAT3 under cytokine stimulation and JAK inhibition (Figure 1). Panel A shows selected cytokine-dose distributions for IFN-*γ*-induced pY-STAT1 and IL-10-induced pY-STAT3, together with the combined full-stimulation reference used in the inhibition experiments. Panel B defines the non-stimulated and fully stimulated reference distributions used to set state thresholds. Panel C shows how inhibitor-treated distributions are redistributed by increasing concentrations of cerdulatinib, and Panel D summarizes the same redistribution as state occupancy. Responses are plotted on the log-response axis, on which the breadth and overlap of the distributions are most readily visualized.

**Figure 1.**
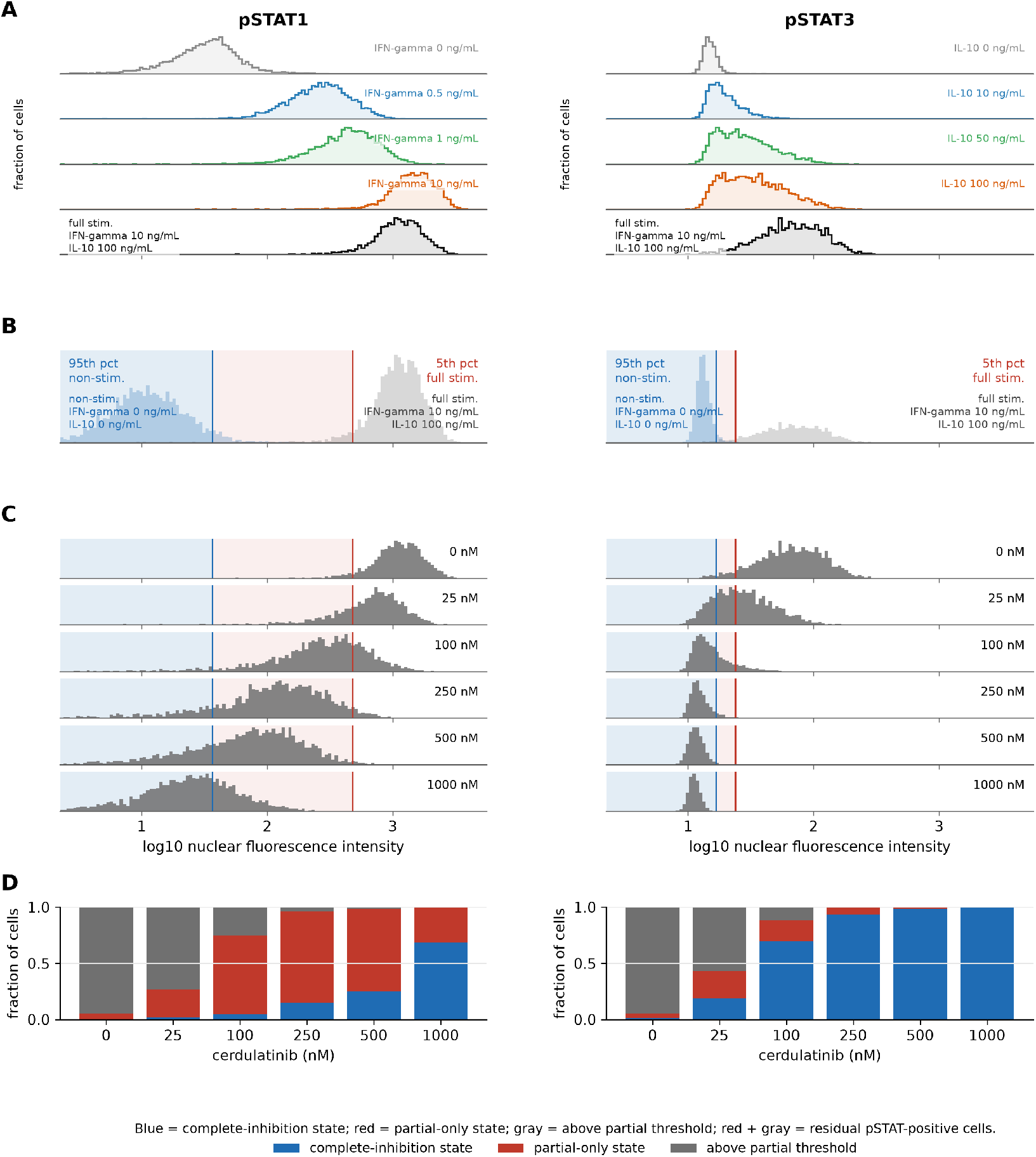
Single-cell response distributions define reference states for inhibitor interpretation. (A) Nuclear pY-STAT1 distributions after IFN-*γ* stimulation and pY-STAT3 distributions after IL-10 stimulation. The final row shows combined full stimulation (IFN-*γ* 10 ng/mL; IL-10 100 ng/mL). Rows are independently y-scaled. (B) Reference distributions: the 95th percentile of the non-stimulated distribution defines the blue complete-inhibition range; the 5th percentile of the fully stimulated distribution defines the upper bound of the red partial-only range. (C) Responses to increasing cerdulatinib doses after full stimulation, relative to the fixed thresholds in B. (D) Corresponding occupancy: blue, complete inhibition; red, partial-only inhibition; gray, above the partial threshold. Red plus gray constitutes the residual pY-STAT-positive fraction. Measurements are pooled across available technical replicates of one biological replicate; a second independent replicate showed similar qualitative patterns.

Two features of these distributions motivate the subsequent analysis. First, the distributions are broad and overlapping: cells exposed to different cytokine concentrations can exhibit similar responses, and even under strong stimulation a fraction of cells remains close to the low-response range. Stimulation therefore redistributes cells across a broad range of responses rather than simply shifting a single characteristic response. Second, the form of this redistribution differs between the two readouts. For pY-STAT1, increasing IFN-*γ* concentration produces a predominantly graded response, with the distribution shifting progressively toward higher values. For pY-STAT3, increasing IL-10 concentration broadens and flattens the distribution over the examined range, indicating that dose affects both the fraction of cells with elevated pY-STAT3 and the response magnitude among those cells. These patterns show that a population summary alone cannot specify how inhibitor treatment redistributes cells among distinct response ranges.

Panels B–D introduce the state-based inhibition metrics used throughout the manuscript. The complete-inhibition threshold is defined by the 95th percentile of the non-stimulated reference distribution. Cells below this threshold are assigned to the complete-inhibition state. The partial-inhibition threshold is defined by the 5th percentile of the fully stimulated reference distribution, constrained not to fall below the complete threshold when the references overlap (Methods). The partial-inhibition metric is the fraction of cells below this threshold and therefore includes both cells in the complete-inhibition state and cells in the intermediate range between the two thresholds. In Figure 1, blue denotes the complete-inhibition range, red the intermediate partial-only range, and gray responses above the partial-inhibition threshold. Red and gray together constitute the residual pY-STAT-positive fraction.

The cerdulatinib-treated distributions show increasing occupancy of the intermediate partial-only range as concentration rises, followed by greater occupancy of the non-stimulated range at higher doses. These are comparisons of cell populations at a fixed endpoint, not trajectories of the same cells. The histograms and stacked occupancy bars show why strong attenuation of a population-level response can coexist with limited occupancy of the non-stimulated range.

### Representative inhibition curves separate population suppression from complete-inhibition state occupancy

Representative THP-1 dose-response profiles illustrate why population suppression and state occupancy cannot be treated as equivalent pharmacological endpoints (Figure 2). Each panel overlays arithmetic-mean inhibition, log-mean inhibition, partial-inhibition state occupancy, and complete-inhibition state occupancy. Solcitinib and cerdulatinib were selected because together they show the principal relationships among these endpoints for both pY-STAT1 and pY-STAT3; the corresponding curves for all inhibitors, cell lines, and readouts are shown in Supplementary Figure S1.

**Figure 2.**
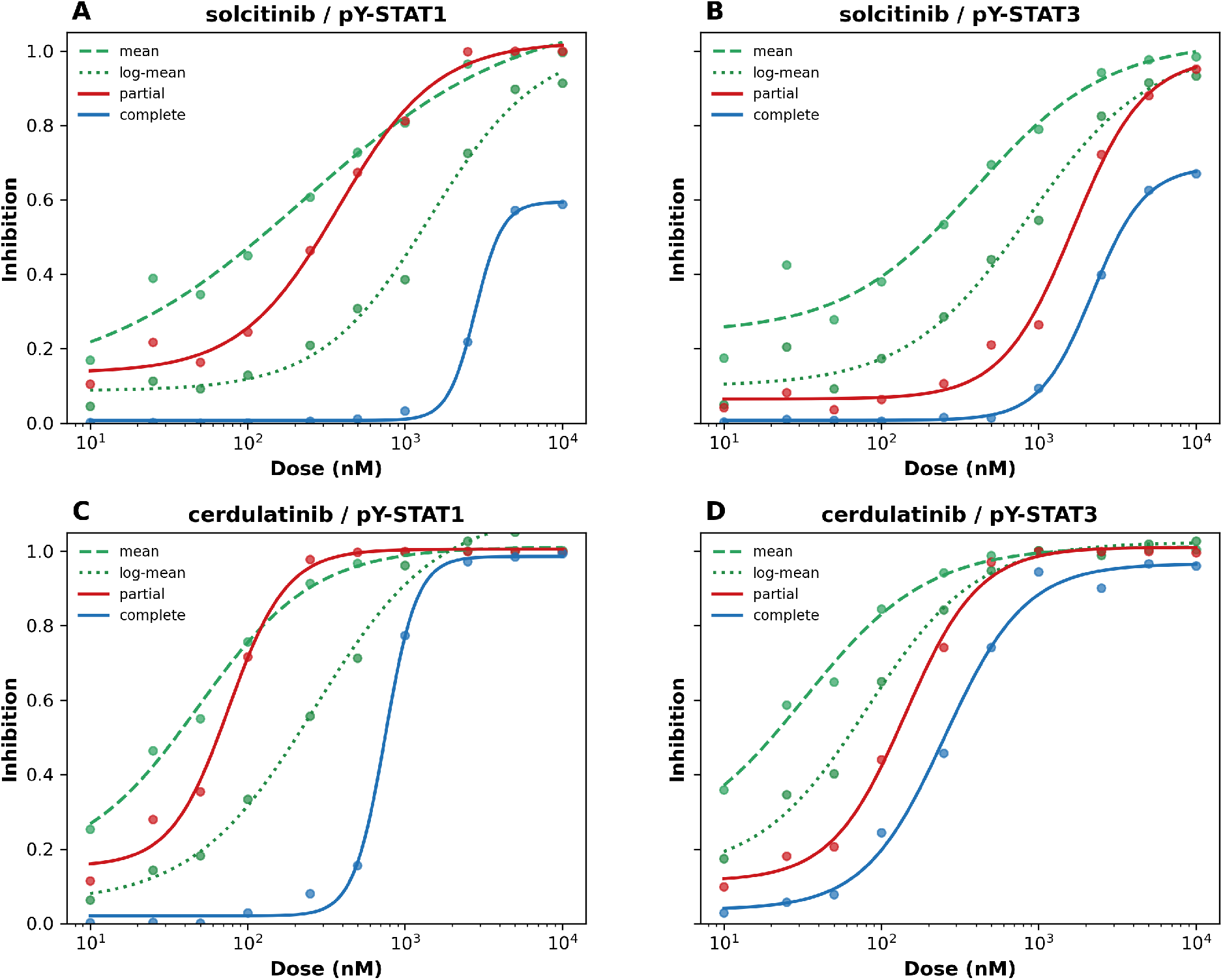
Representative inhibition curves reveal dissociation between population suppression and state occupancy. Representative dose-response curves in THP-1 cells for solcitinib (A, B) and cerdulatinib (C, D), measured using pY-STAT1 (A, C) and pY-STAT3 (B, D) as readouts after cytokine stimulation. Each panel shows four inhibition endpoints: arithmetic-mean inhibition, log-mean inhibition, partial-inhibition state occupancy, and complete-inhibition state occupancy. Points represent observed values at each inhibitor dose; lines show fitted dose-response curves. Arithmetic-mean inhibition provides the most left-shifted estimate of inhibition, log-mean inhibition shifts the population endpoint to higher doses, and complete-inhibition state occupancy was generally the most right-shifted endpoint. Points represent pooled observations from two technical replicates of one representative biological replicate, and fitted curves were generated from these pooled data; similar qualitative relationships were observed in a second independent biological replicate.

Across all four panels, arithmetic-mean inhibition was left-shifted relative to log-mean inhibition, indicating lower apparent inhibitory concentrations when responses were averaged on the raw scale. Complete-inhibition state occupancy was generally displaced furthest to the right. Partial-inhibition state occupancy remained closer to the population summaries than complete-inhibition state occupancy, consistent with all three endpoints being influenced by the bulk of the responding distribution. Thus, the endpoints did not follow a single fixed ordering in every condition, but they consistently showed that attenuation of the typical response and entry into the complete-inhibition state can occur at different inhibitor concentrations.

This separation is clearest for cerdulatinib-inhibited pY-STAT1 (Figure 2C). At intermediate concentrations, arithmetic-mean inhibition is close to saturation, and both log-mean inhibition and partial-inhibition state occupancy are high, yet complete-inhibition state occupancy remains low. The population response can therefore be strongly attenuated while many cells remain above the non-stimulated range. A similar separation is visible for solcitinib-inhibited pY-STAT3 (Figure 2B), where both population summaries indicate substantial inhibition before complete-inhibition state occupancy rises appreciably.

These examples show that the discrepancy is not restricted to the known difference between arithmetic averaging and analysis on the log-response scale. The log-mean provides a less left-shifted population benchmark than the arithmetic mean, but it still does not specify how many cells have entered the non-stimulated response range. The following analysis tests how consistently this separation occurs across the full inhibitor–cell-line–readout panel.

### Matching population inhibition to state occupancy reveals residual pY-STAT-positive fractions

We next used the full inhibitor–cell-line–readout panel to establish the empirical relationship between population inhibition and single-cell state occupancy. Across 24 profiles, Figure 3 asks what fractions of cells had left the fully stimulated response range or entered the complete-inhibition state at matched levels of arithmetic-mean or log-mean inhibition. This comparison quantifies both how informative a population-inhibition curve is about the underlying redistribution of single-cell responses and how large a residual pY-STAT-positive fraction may remain.

**Figure 3.**
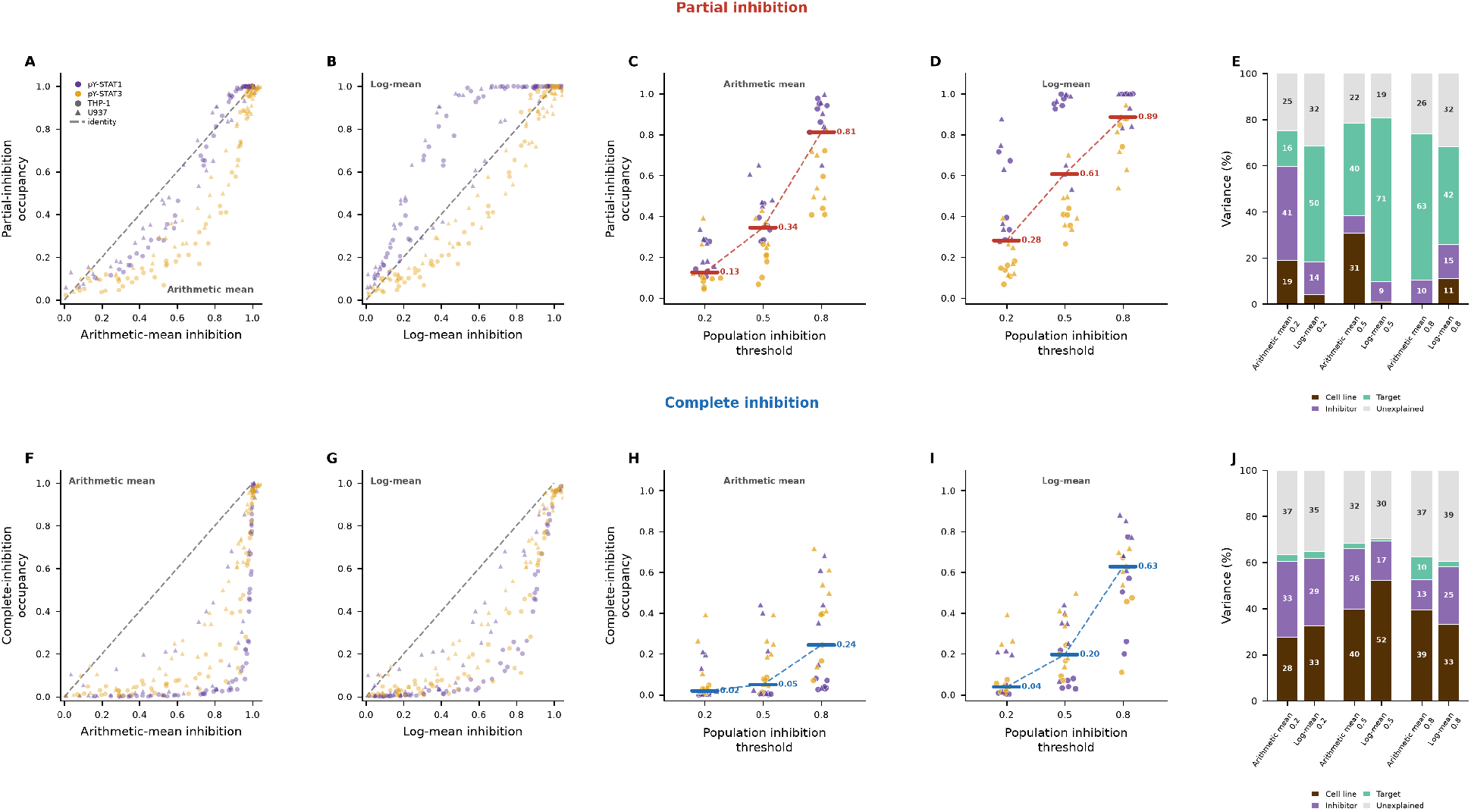
Matching population inhibition to state occupancy quantifies residual pY-STAT-positive fractions. Partial-inhibition occupancy (A–E) is the fraction of cells below the fully stimulated response range; complete-inhibition occupancy (F–J) is the fraction within the non-stimulated response range. Violet and amber denote pY-STAT1 and pY-STAT3; circles and triangles denote THP-1 and U937. (A, B) Partial-inhibition occupancy versus arithmetic-mean inhibition (A) or log-mean inhibition (B) at the same measured doses. Dashed line, identity. (C, D) Partial-inhibition occupancy at the lowest measured dose at which the fitted population-summary curve reached or exceeded 0.2, 0.5, or 0.8. Each point represents one profile; colored bars show medians and dashed lines connect them. (E) Descriptive variance shares associated with cell line, inhibitor, phospho-target, and an unexplained remainder for each benchmark and population summary (Methods). (F, G) Complete-inhibition occupancy versus arithmetic-mean inhibition (F) or log-mean inhibition (G) at the same measured doses. (H, I) Complete-inhibition occupancy using the threshold-crossing procedure in C and D. At the 0.8 benchmark, median occupancy was 0.24 for arithmetic-mean inhibition and 0.63 for log-mean inhibition, leaving median residual pY-STAT-positive fractions of 76% and 37%, respectively. (J) Between-profile variance partitioned as in E. Cell line, inhibitor, and phospho-target together accounted for approximately 60–70% of the variance; cell line was the largest identifiable contributor in most configurations and on average (label-permutation test, *p* ≤ 0.003). *n* = 22–24 evaluable profiles per benchmark; profiles whose fitted curve did not reach a benchmark were omitted.

Partial-inhibition occupancy remained relatively close to both population summaries across doses (Figure 3A,B). At the lowest measured dose at which fitted log-mean inhibition reached each benchmark (0.2, 0.5, or 0.8), median partial-inhibition occupancy was 0.28, 0.61, and 0.89, respectively (Figure 3C,D). The log-mean therefore provided a close, although not exact, benchmark for the fraction of cells that had left the fully stimulated response range. The remaining profile-to-profile differences were structured: variance partitioning identified phospho-target as the largest identifiable contributor in most threshold–summary combinations and on average (Figure 3E).

Complete-inhibition occupancy was more loosely related to population inhibition. Across the same profiles, the fraction of cells entering the non-stimulated response range was generally lower than the corresponding population-summary inhibition (Figure 3F,G). Applying the same lowest-dose criterion across inhibition benchmarks revealed a nonlinear relationship. At arithmetic-mean inhibition levels of 0.2, 0.5, and 0.8, median complete-inhibition occupancy was 0.02, 0.05, and 0.24, respectively. At the corresponding log-mean inhibition levels, median occupancy was 0.04, 0.20, and 0.63 (Figure 3H,I). Thus, the log-mean approached complete-inhibition occupancy more closely at high levels of population suppression than did the arithmetic mean. Even at the 0.8 benchmark, however, the discrepancy remained substantial across individual profiles. At the lowest measured dose at which arithmetic-mean inhibition reached 0.8, complete-inhibition occupancy had an interquartile range of 0.07–0.47 and remained below 0.3 in 12 of 23 evaluable profiles. At the corresponding log-mean benchmark, complete-inhibition occupancy remained below 0.5 in 5 of 22 evaluable profiles. Expressed as residual fractions, doses meeting the 80% arithmetic-mean or log-mean inhibition benchmark still left median residual pY-STAT-positive fractions of 76% and 37%, respectively. The log-mean therefore provided a useful quantitative expectation, but neither population summary uniquely specified the fraction of cells that had entered the non-stimulated response range.

The residual pY-STAT-positive fraction at matched population inhibition also differed systematically among profiles. Cell line, inhibitor, and phospho-target together accounted for approximately 60–70% of the between-profile variance in complete-inhibition occupancy at the examined thresholds. Cell line provided the largest identifiable contribution in most configurations and on average (label-permutation test on cell line, *p* ≤ 0.003; Figure 3J), whereas 30–40% of the variance was not explained by the factors included in the analysis. In contrast, the largest identifiable contribution to variation in partial-inhibition occupancy was generally associated with phospho-target rather than cell line (Figure 3E). These are descriptive associations within this panel: only two cell lines were examined, and batch is confounded with cell line and inhibitor panel (Methods).

Together, our data provide an empirical guide to the cell fractions expected from a conventional inhibition curve. Within this panel, log-mean inhibition approximates departure from the stimulated range and helps estimate complete-inhibition occupancy, including the variation among conditions. The relationship is informative but not deterministic: direct occupancy measurements remain necessary to establish the fraction above the non-stimulated pY-STAT range in a particular condition. The next analysis expresses this separation as a gap in apparent IC_50_.

### Complete-inhibition occupancy shows a median ∼2.4-to 7.4-fold dose gap from population summaries

The distinction between population inhibition and state occupancy has a direct pharmacological consequence: the same inhibitor profile can yield different apparent potencies depending on the endpoint. We quantified these differences across all 24 profiles by comparing the apparent IC_50_ values of arithmetic-mean and log-mean inhibition with those of partial- and complete-inhibition occupancy (Figure 4).

**Figure 4.**
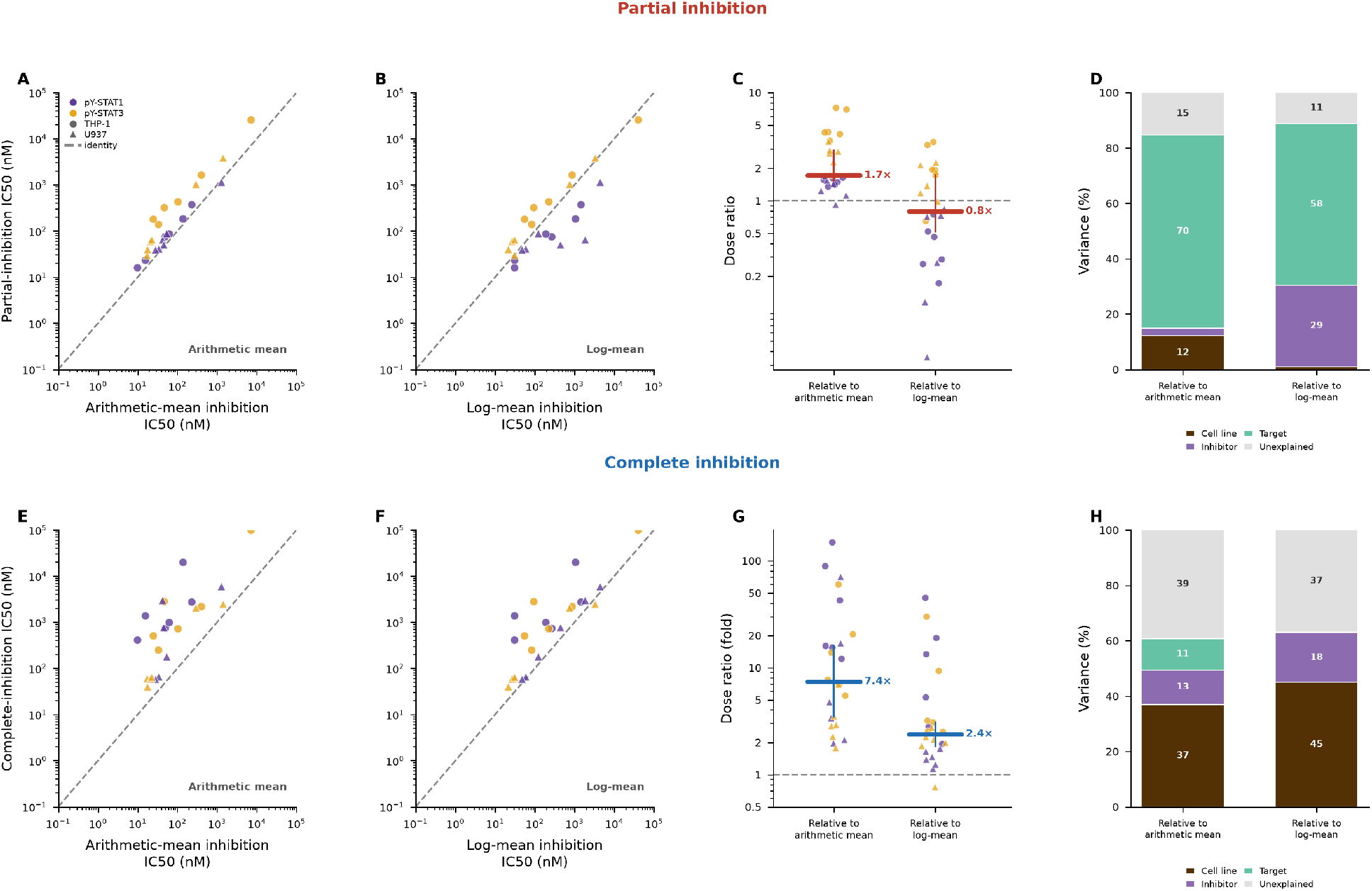
Partial-inhibition IC_50_ approximates log-mean IC_50_, whereas complete-inhibition IC_50_ is right-shifted. The figure compares partial-inhibition occupancy (A–D) and complete-inhibition occupancy (E–H) with arithmetic-mean and log-mean inhibition across 24 inhibitor–cell-line–readout profiles. Violet and amber denote pY-STAT1 and pY-STAT3; circles and triangles denote THP-1 and U937. (A, B) Apparent partial-inhibition IC_50_ versus arithmetic-mean IC_50_ (A) or log-mean IC_50_ (B). Both axes are logarithmic; the dashed line denotes identity. (C) Per-profile ratios of partial-inhibition IC_50_ to each population-summary IC_50_. Points show individual profiles, the horizontal colored bar denotes the median, the vertical line denotes its 95% confidence interval obtained by bootstrap resampling of the 24 profiles (10,000 resamples), and the dashed line denotes a ratio of one. Median ratios were 0.8 relative to the log-mean (95% confidence interval, 0.5–1.7) and 1.7 relative to the arithmetic mean (95% confidence interval, 1.5–2.9). (D) Between-profile variance in the log10-transformed ratios from C, partitioned among cell line, inhibitor, phospho-target, and an unexplained remainder. (E, F) Apparent complete-inhibition IC_50_ versus arithmetic-mean IC_50_ (E) or log-mean IC_50_ (F), plotted as in A and B. (G) Per-profile ratios plotted as in C. Median ratios were 2.4 relative to the log-mean (95% confidence interval, 1.9–3.1) and 7.4 relative to the arithmetic mean (95% confidence interval, 3.5–16). (H) Variance in log10-transformed ratios from G, partitioned as in D. Factor shares are descriptive and do not separate cell line from batch; confidence intervals do not represent biological-replicate uncertainty (Methods).

Partial-inhibition IC_50_ values lay close to the population-summary estimates, particularly the log-mean (Figure 4A,B). The median ratio of partial-inhibition IC_50_ to log-mean IC_50_ was 0.8 (95% confidence interval, 0.5–1.7), compared with 1.7 relative to the arithmetic-mean IC_50_ (95% confidence interval, 1.5–2.9; Figure 4C). Thus, the log-mean provided a close estimate of the concentration range over which cells left the stimulated response range.

Complete-inhibition IC_50_ values showed a clear rightward displacement from the population-summary estimates (Figure 4E,F). The complete-inhibition IC_50_ exceeded the log-mean IC_50_ in 23 of 24 profiles. The median fold difference was 2.4 relative to the log-mean (95% confidence interval, 1.9–3.1) and 7.4 relative to the arithmetic mean (95% confidence interval, 3.5–16; Figure 4G). Moving from arithmetic-mean to log-mean inhibition therefore reduced the dose gap substantially, but the fitted midpoint of complete-inhibition occupancy remained at higher concentrations than the log-mean midpoint.

The fold differences varied among profiles. Phospho-target was the largest identifiable contributor to variation in the partial-inhibition ratios (Figure 4D). For complete-inhibition ratios, cell line was the largest identifiable contributor, accounting for 37–45% of the between-profile variance depending on the population-summary comparator. Inhibitor made a smaller contribution, while 37–39% of the variance remained unexplained by the three included factors (Figure 4H). The cell-line association is specific to this panel and cannot be separated from batch.

Together, population-summary IC_50_ values, particularly the log-mean, approximated partial-inhibition IC_50_ but generally underestimated complete-inhibition IC_50_. Apparent potency therefore depends on the pharmacological endpoint: attenuation of the typical response and entry of cells into the non-stimulated response range are separated by a measurable, condition-dependent dose gap.

## Discussion

The central contribution of this study is a quantitative link between conventional inhibitor-response curves and the redistribution of single-cell pY-STAT1/3 responses. Within the tested panel, our data help estimate the fractions of cells that have left the stimulated response range or entered the non-stimulated range at a given level of population suppression. The measured dose gap shows why an apparent potency based on the population average need not locate the concentration regime in which individual cells return to baseline. Its magnitude and variability matter when the experimental aim is to reduce the fraction of cells retaining elevated pY-STAT responses, rather than only lower their average signal.

This distinction is meaningful because cytokine-response heterogeneity can reflect structured differences in molecular content, signaling capacity, and cellular state rather than simply a loss of signaling fidelity [5, 18–20]. The same average can arise from distributions with different shapes and minority-cell fractions, with different biological implications [3, 4]. Information may therefore be carried by the fractions of cells occupying response ranges as well as by shifts in central tendency [6, 21, 22]. Partial inhibition measures departure from the stimulated range, whereas complete inhibition measures entry into the non-stimulated range. Neither fraction is interchangeable with an average.

Previous studies established that potency alone can conceal different modes of drug response and that threshold-based single-cell measurements resolve features hidden by population summaries [8– 10]. Our results quantify these relationships across a pharmacologically diverse JAK-inhibitor panel. The log-mean provides a scale-appropriate population benchmark for these broad distributions and generally approximates departure from the stimulated range. Yet the apparent IC_50_ for complete-inhibition occupancy remains a median 2.4-fold higher than the log-mean IC_50_, compared with 7.4-fold relative to the arithmetic mean. Changing the averaging scale reduces, but does not remove, the dose gap.

The biological relevance lies in the possibility that cells retaining elevated responses can influence outcomes disproportionate to their abundance. Heterogeneous subpopulations have been associated in other systems with divergent cell-fate decisions, inflammatory responses, and persistence under treatment [7, 23–25]. Such consequences remain plausible rather than demonstrated here. Our data identify an upstream response regime in which they could matter: strong average suppression accompanied by cells remaining above the non-stimulated pY-STAT range. Whether those cells sustain distinct transcriptional programs, paracrine effects, or survival decisions requires direct functional testing.

STAT complex formation provides a possible biological context for the response variability. Canonical activated dimers are stabilized by reciprocal phosphotyrosine–SH2 interactions [26]. In STAT1, N-terminal-domain-dependent dimer reorientation also permits access to dephosphorylation [27]. The balance between STAT1 and STAT3 homo- and heterodimeric complexes can affect transcription: in a type-I-interferon setting, STAT3 reduced DNA-binding STAT1 homodimers and STAT1-dependent gene activation without reducing STAT1 tyrosine phosphorylation or nuclear translocation [28]. Variation in STAT abundance and phosphorylation turnover could therefore influence both the persistence and functional consequences of nuclear pY-STAT responses. Our measurements, however, resolve tyrosine phosphorylation rather than dimer composition; they establish neither a dimerization mechanism for the dose gap nor the transcriptional activity of cells remaining above the non-stimulated range.

The two occupancy endpoints differed in the factors associated with their variability. Phospho-target was most strongly associated with variation in partial inhibition, whereas cell line provided the largest identifiable contribution for complete inhibition and its apparent IC_50_ shift within this panel. These associations do not identify a mechanism, particularly because batch is confounded with cell line and inhibitor panel. Approximately 40% of the variation in the complete-inhibition dose ratio remained unexplained by the three factors; this remainder also includes omitted interactions, measurement error, and fit uncertainty, not only cellular heterogeneity. The correspondence between partial inhibition and log-mean inhibition is likewise empirical, not a mathematical identity.

Several limitations define the scope of these conclusions. First, the study does not identify the molecular determinants of the remaining pY-STAT-positive fraction. Receptor abundance, pathway-component levels, and cellular state are plausible contributors, but were not measured directly [4, 5, 19].

Second, partial and complete inhibition are operational definitions derived from reference distributions, not uniquely privileged biological thresholds. “Complete inhibition” means that nuclear pY-STAT1 or pY-STAT3 falls within the non-stimulated range at the measured endpoint. It does not establish complete silencing of JAK-STAT signaling: other STAT proteins, earlier or later activity, downstream transcription, and responses to subsequent stimulation were not measured [29]. Conversely, a response above the threshold is not by itself proof of a distinct functional fate.

Third, apparent IC_50_ estimates are fitted midpoints between plateaus, not doses that place every cell in the complete-inhibition range. Estimates for this occupancy endpoint are particularly sensitive to incomplete saturation and limited dynamic range. The fold differences are therefore comparative summaries, not universal dose corrections. Their direction persisted in the second biological replicate, supporting the qualitative result; two biological replicates do not establish a broadly generalizable numerical gap. Bootstrap intervals across profiles describe the sampled panel rather than uncertainty across independent biological replicates.

Finally, the panel comprises two PMA-differentiated cell lines, two pY-STAT readouts, six inhibitors, and one post-stimulation time point. It does not establish cell line as a generally dominant determinant or distinguish pathway-selective inhibition from other drug effects. Additional cell types, particularly primary cells, and time points would be needed to establish how broadly the quantitative relationships apply.

Together, these results provide an empirical guide to the cell fractions expected at a given level of population inhibition, including the variation that limits such estimates. Single-cell inhibitor profiling should report both scale-appropriate population summaries and state-occupancy endpoints. For heterogeneous signaling systems, attenuation of the typical response and entry of individual cells into the non-stimulated pY-STAT range are distinct pharmacological outcomes, and the dose gap between them warrants measurement in its own right.

## Methods

### Study design

Single-cell imaging experiments quantified JAK-STAT inhibition in two human monocytic macrophage-like cell lines, THP-1 and U937, using nuclear pY-STAT1 and pY-STAT3 as readouts of cytokine response. The study comprised four dose-response experiments spanning six JAK-pathway inhibitors: solcitinib, BMS-911543, cerdulatinib, baricitinib, ruxolitinib, and tofacitinib. This panel was analyzed as a set of pharmacologically diverse perturbations of shared JAK-family signaling, not as a biochemical selectivity screen. Across the combined dataset, inhibitor effects were quantified using four inhibition endpoints: arithmetic mean, log-mean, partial inhibition, and complete inhibition. The integrated dataset comprised 24 inhibitor-cell-line-readout profiles and 96 fitted dose-response curves across the four inhibition metrics. Experiments were performed in two independent biological replicates, each comprising two technical replicates per condition. The figures shown are based on pooled single-cell measurements from available technical replicates after filtering in one representative biological replicate. To verify reproducibility, we processed the second biological replicate of each of the four experiments through the identical pipeline and compared the resulting IC_50_ values with those of the shown replicate (Supplementary Figure S2).

### Cell culture, macrophage-like differentiation, and cytokine stimulation

THP-1 (TIB-202, ATCC) and U937 (CRL-1593.2, ATCC) cells were cultured under standard conditions at 37 °C in a humidified atmosphere of 5% CO_2_/95% air. Cells were maintained in RPMI 1640 medium (Corning, #10-040-CV) supplemented with 10% fetal bovine serum (ThermoFisher, #10500064) and 1% penicillin-streptomycin (ThermoFisher, #15140122); THP-1 medium was additionally supplemented with 0.05 mM 2-mercaptoethanol (Gibco, #21985023). For macrophage-like differentiation, THP-1 and U937 cells were suspended in medium containing 20 ng/mL phorbol 12-myristate 13-acetate (PMA; Sigma-Aldrich, #P1585) and plated in 96-well *µ*Clear flat-bottom imaging plates (Greiner, #655090) at 2 *×* 10^4^ cells per well. After 24 h, PMA-containing medium was removed and replaced with fresh medium, and cells were maintained for an additional 72 h before inhibitor treatment.

For the original inhibitor panel, differentiated THP-1 and U937 cells were incubated for 2 h with solcitinib (Selleckchem, #S5917), BMS-911543 (Selleckchem, #S7144), or cerdulatinib (Selleckchem, #S7634) at ten concentrations spanning 10–10,000 nM before cytokine stimulation. Cells were then stimulated for 30 min with recombinant human IFN-*γ* (10 ng/mL; ThermoFisher, #PHC4031) and recombinant human IL-10 (100 ng/mL; PeproTech, #200-10), which served as the full-stimulation condition in the inhibition experiments.

For the additional clinically used inhibitors, differentiated THP-1 and U937 cells were incubated with baricitinib (Selleckchem, #S2851), ruxolitinib (Selleckchem, #S1378), or tofacitinib (Selleckchem, #S2789) at ten concentrations spanning 10–10,000 nM for 2 h before cytokine stimulation; in the U937 experiment baricitinib was additionally applied at 0.5 nM. Cells were then stimulated under the same full-stimulation conditions as above.

### Immunofluorescence imaging and single-cell signal extraction

After stimulation, cells were fixed with 3.7% paraformaldehyde (Sigma-Aldrich, #P6148) for 10 min at room temperature, permeabilized with 90% ice-cold methanol (Sigma, #322415) for 30 min at −20 °C, and blocked in 5% bovine serum albumin (Merck, #821006) containing 0.3% Triton X-100 (Sigma-Aldrich, #T9284) for 1 h at room temperature. Cells were then incubated overnight at 4 °C with anti-phospho-STAT1 (Tyr701) antibody (Cell Signaling, #9167; 1:100) and/or anti-phospho-STAT3 (Tyr705) antibody (Cell Signaling, #4113; 1:200) diluted in 1% bovine serum albumin with 0.3% Triton X-100. The following day, cells were incubated with Alexa Fluor-conjugated secondary antibodies (ThermoFisher, #A-21206; 1:500 and ThermoFisher, #A-31570; 1:500) for 1.5 h at room temperature and counterstained with 2 *µ*g/mL DAPI (Sigma-Aldrich, #D9542) for 10 min.

Fluorescence images were acquired using an automated confocal microscope (Pathway 435, BD; objective UAPO/340 20*×*, numerical aperture 0.75, Olympus) and analyzed using the original image-analysis workflow based on CellProfiler, ImageJ, and custom scripts. Nuclear fluorescence intensities were quantified at single-cell resolution. Signal extraction was based on the mean fluorescence intensity of nuclear pixels after background correction. The analysis pipeline rescaled exported intensities by 65,535 and retained finite values strictly between 1 and 10^4^ counts, separately for each readout. Groups and reference distributions required at least 10 retained nuclei. Well exclusions and their provenance are recorded in the deposited metadata. Available cells from technical replicates were pooled after filtering; the U937 newer-inhibitor experiment retained one non-stimulated reference well, and the THP-1 original-inhibitor experiment retained one fully stimulated reference well.

### Response representation and inhibition metrics

Single-cell signaling responses were represented and analyzed on the log-response axis. In the text, pY-STAT mean refers to the arithmetic mean single-cell tyrosine-phosphorylation signal within a condition, measured as nuclear phospho-STAT fluorescence intensity for pY-STAT1 or pY-STAT3. Log-mean was defined as the arithmetic mean of the corresponding log10-transformed single-cell signal. Following quality-control filtering, all analyzed signal values were strictly positive, so the log transformation was applied without a pseudocount.

Arithmetic-mean inhibition was defined from the condition-level arithmetic mean of the raw single-cell response. For each experiment and readout, the no-inhibitor fully stimulated reference was set to 0 inhibition and the no-inhibitor non-stimulated reference was set to 1 inhibition:

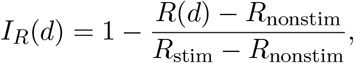

where *R*(*d*) is the response summary at inhibitor dose *d, R*_nonstim_ is the corresponding summary of the no-inhibitor non-stimulated reference, and *R*_stim_ is the corresponding summary of the no-inhibitor fully stimulated reference. Log-mean inhibition was calculated by applying the same normalization to the mean of log10-transformed single-cell responses. Thus, mean-based inhibition metrics and complete-inhibition state occupancy were anchored to the same non-stimulated reference distribution, but used it differently: mean and log-mean inhibition used its central tendency as the low-signal normalization endpoint, whereas complete inhibition used its 95th percentile as the threshold for the complete-inhibition state.

Complete and partial inhibition were defined from percentile thresholds of the reference distributions. The complete-inhibition threshold was the 95th percentile of the non-stimulated reference distribution. The partial-inhibition threshold was the 5th percentile of the fully stimulated reference distribution, constrained not to fall below the complete threshold when the reference distributions overlapped. Complete and partial inhibition were the fractions of retained cells strictly below their respective thresholds. These fractions were not baseline-corrected; the non-stimulated reference itself therefore has approximately 95% complete-inhibition occupancy. In the U937 pY-STAT3 profiles for baricitinib, ruxolitinib, and tofacitinib, overlap of the reference tails made the thresholds coincide, so the partial- and complete-inhibition curves overlap (Supplementary Figure S1).

### Dose-response fitting and apparent IC50 estimation

For each combination of inhibitor, cell line, readout, and inhibition metric, dose-response curves were fit with a four-parameter Hill-type function:

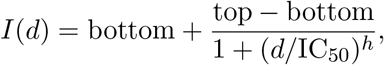

where *I*(*d*) denotes the inhibition endpoint at inhibitor dose *d*, and bottom, top, IC_50_, and Hill slope *h* were treated as free parameters. With *h >* 0 this function returns ‘top’ as *d* → 0 and ‘bottom’ as *d* → ∞; because inhibition increases with dose, fitted profiles have bottom *>* top, and the curve rises. The fitted IC_50_ is therefore the dose at which *I*(*d*) lies halfway between the two fitted plateaus, not necessarily the dose at which *I*(*d*) = 0.5. No-inhibitor controls defined the reference distributions; only positive inhibitor doses were included in the dose-response fits.

Fits were performed in Python using `scipy.optimize.curve_fit`. Initial parameter values were fixed at bottom_0_ = 0, top_0_ = 1, IC_500_ = median of the nonzero doses, and *h*_0_ = 1. Parameter bounds were imposed as follows: bottom and top were constrained to [0, 1.2], IC_50_ to [minimum nonzero dose/10, maximum dose *×* 10], and *h* to [0.2, 5.0].

All 96 curves (24 profiles *×* 4 metrics) converged, so every IC_50_ reported here is a fitted midpoint and no fallback estimator was applied; a curve with fewer than four nonzero dose levels or a failed fit would have been excluded rather than mixed with the fitted values. The IC_50_ bound permitted estimates up to ten times the highest tested concentration, so some estimates are extrapolated beyond the measured dose window: of the 96 fits, four exceeded the maximum tested concentration of 10,000 nM (one log-mean, one partial, and two complete), and these fall in just two of the 24 profiles, both BMS-911543 in THP-1 cells; one of the two complete fits reached the upper fitting bound. Twenty-two of the 24 profiles have all four IC_50_ values bracketed by tested concentrations; restricting the analysis to those profiles left the reported medians essentially unchanged (complete over log-mean 2.2 rather than 2.4, 95% confidence interval 1.8–3.0; complete over arithmetic mean 7.1 rather than 7.4, 95% confidence interval 3.4–16), so the dose gap is not an artifact of extrapolated estimates. Per-profile fit status, plateau values and bracketing are given in the deposited source table.

### Integrated comparison across conditions

Condition-level IC_50_ comparisons were descriptive and comparative. The principal comparisons used in Figure 4 were IC_50_(complete) versus IC_50_(log-mean) and IC_50_(complete) versus IC_50_(mean). Scatter plots were shown on log-log axes with an identity line indicating equal potency under the two compared inhibition definitions, with points encoded by phospho-target (color) and cell line (marker shape).

To summarize the direction and magnitude of these shifts, we computed, for each profile, the ratio of the fitted complete-inhibition midpoint to each population-summary midpoint:

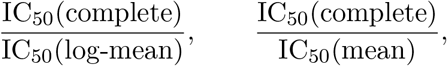

We report the median across profiles with a 95% confidence interval obtained by bootstrap resampling of profiles (10,000 resamples). The ratios compare endpoint-specific midpoints, not concentrations required to suppress every cell. The intervals describe resampling of this panel; profiles share experiments and controls, and the intervals do not quantify between-biological-replicate uncertainty or propagate individual fit uncertainty. To describe associations with profile-to-profile variability (Figure 4H), log-transformed fold-ratios were partitioned across cell line, inhibitor, and phospho-target using the same three-factor type-II analysis of variance as for state occupancy.

### Threshold-crossing and variance-decomposition analysis

To relate population-level suppression to single-cell state occupancy (Figure 3), we identified, for each profile, the lowest tested dose at which the fitted arithmetic-mean or log-mean inhibition curve reached or exceeded a target level (0.2, 0.5, or 0.8). We recorded the observed partial- and complete-inhibition fractions at that dose, without interpolating the occupancy between doses. Profiles not reaching a target within the tested range were omitted at that target. Medians were summarized with profile-bootstrap 95% confidence intervals (10,000 resamples), subject to the limitations above.

Between-profile variability in both occupancy endpoints was summarized by an additive fixed-effects model with cell line, inhibitor, and phospho-target. Each type-II factor sum of squares and the residual sum of squares was divided by their sum and expressed as a percentage. In unbalanced subsets this normalization is not necessarily identical to a partition of the raw total sum of squares; the reported percentages are descriptive model-based shares, not random-effects variance components. With 24 profiles and one observation per factor combination, the full factorial model would be saturated. The additive model has 16 residual degrees of freedom for the full panel, pooling omitted interactions with measurement, threshold, and fit uncertainty; interactions cannot be separated from pure error. Experiment is also confounded with cell line and inhibitor panel. We therefore do not interpret the residual as intrinsic cellular variability or factor shares as causal contributions. Cell-line labels were shuffled across profiles in 2,000 permutations, with *p* estimated as (*k* + 1)*/*(*n* + 1). These tests describe association under profile-label permutation; they do not separate cell-line biology from batch or establish effects across independent biological replicates.

## Software and reproducibility

All analyses and figure generation were performed in Python using the numbered analysis scripts developed for this study. The workflow proceeds from the raw per-nucleus exports through per-dose inhibition tables, four-parameter logistic fits and derived summary tables, to deterministic figure assembly. The public repository contains the raw measurements, plate metadata, source tables, manuscript figure assets, executable scripts and automated tests needed to inspect and rerun the analysis rather than relying on hand-transcribed values.

## Data and code availability

The raw per-nucleus image-analysis exports for all imaging experiments, including the second biological replicate, together with plate metadata, source tables, manuscript figure assets, figure-generation scripts, tests and manuscript files, are available at https://github.com/sysbiosig/jak-stat-dose-gap. The repository supports the full figure set: a single command regenerates every derived table and Figures 1, 2, 3, 4 and S1 from the raw files, and the plotted IC50 source table and replicate raw measurements underlying Supplementary Figure S2 are deposited and documented in the same repository. An automated test suite verifies the regenerated values against those reported here. An archived release with a citable DOI will be deposited before publication.

## Declarations

### Funding

This research was supported by the National Science Center, Poland, under grant number 2020/39/B/NZ2/03259.

### Author contributions

K.E.Z., J.W., C.M.W., and A.B. designed and performed the experiments. P.T. and K.N. analyzed the data. A.B. and T.R. provided the cell lines and contributed expertise in their culture and differentiation. R.M. and M.K. interpreted the results and wrote the manuscript. All authors read and approved the final manuscript.

### Competing interests

The authors declare no competing interests.

## Notes

https://github.com/sysbiosig/jak-stat-dose-gap

### Competing Interest Statement

The authors have declared no competing interest.

