## Supplementary Figures S1 and S2 for "Single-cell profiling reveals a dose gap between population-average and complete pY-STAT1/3 suppression by JAK inhibitors"

### Supplementary figure legends

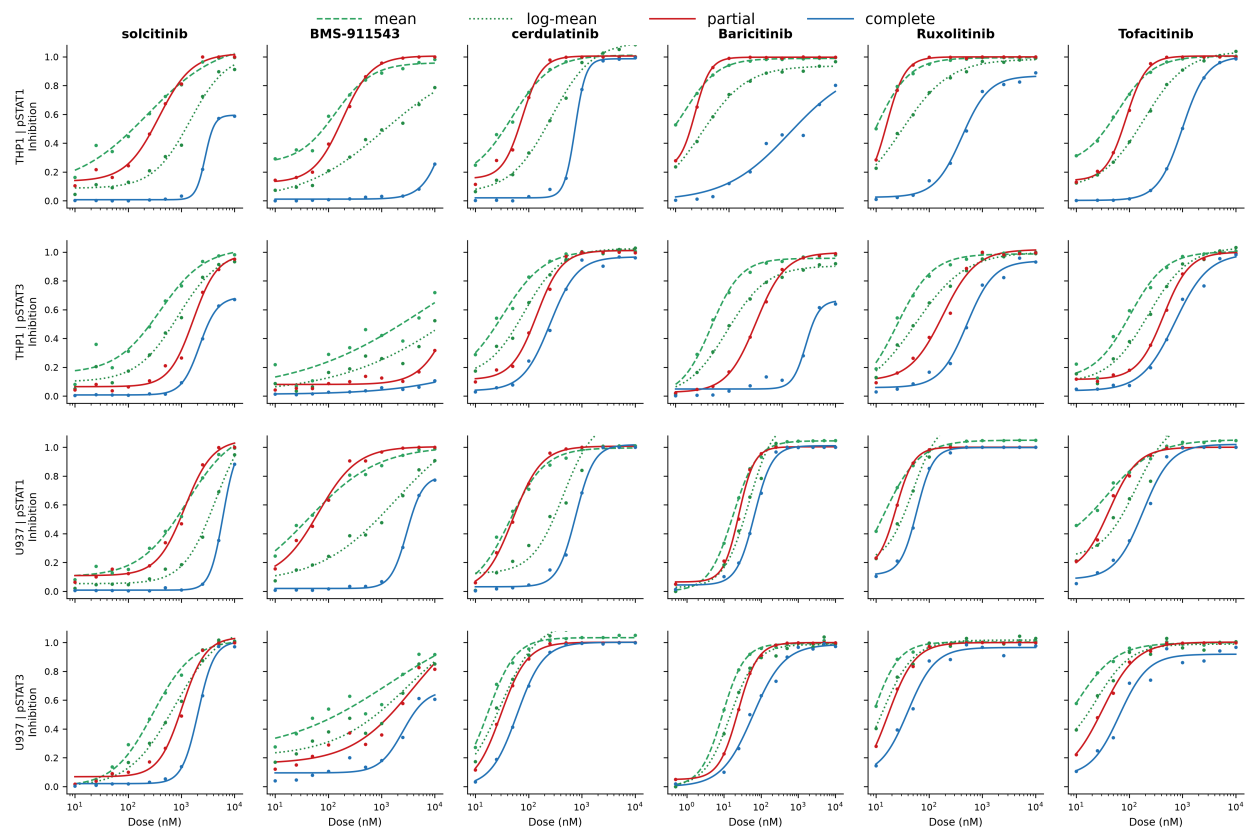

Supplementary Figure S1. **Full dose-response atlas across inhibitors, cell lines, and read-outs.** Observed inhibition values and fitted dose-response curves for arithmetic mean, log-mean, partial inhibition, and complete inhibition across all inhibitor-cell-line-readout combinations analyzed in this study. Columns correspond to the six inhibitors solcitinib, BMS-911543, cerdulatinib, baricitinib, ruxolitinib, and tofacitinib. Rows correspond to THP-1 pY-STAT1, THP-1 pY-STAT3, U937 pY-STAT1, and U937 pY-STAT3.

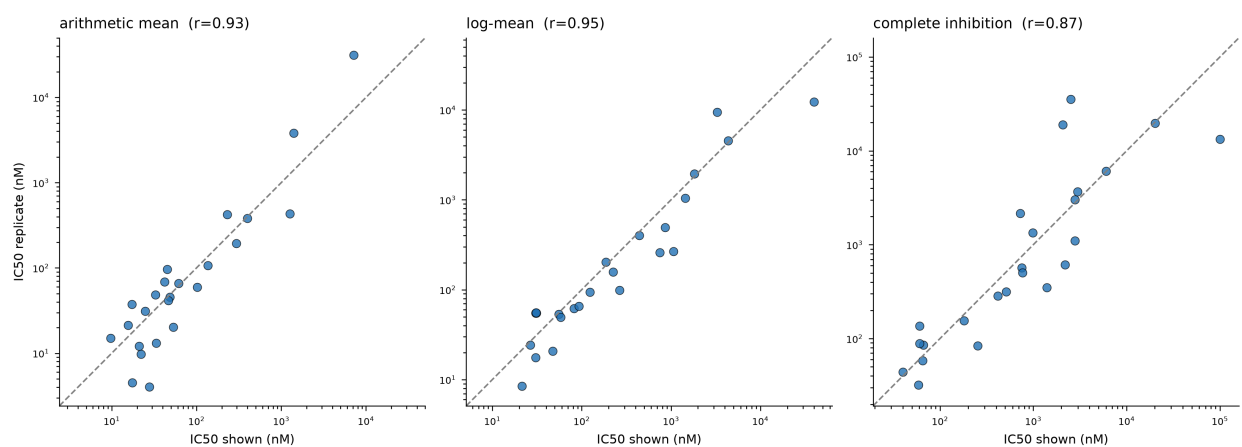

Supplementary Figure S2. **Comparison of apparent IC<sub>50</sub> estimates across biological replicates.** For each of the four dose-response experiments, apparent IC<sub>50</sub> estimates from the deposited concordance table for the second biological replicate were compared with those of the representative replicate shown in the main figures. Each point is one inhibitor-cell-line-readout profile (24 profiles total); axes are shown on a logarithmic scale and the dashed line marks identity. Panels correspond to the arithmetic mean, log-mean, and complete-inhibition metrics; the log-log Pearson correlation is given in each panel title. The relative ordering of the metrics, including IC<sub>50</sub>(complete) exceeding IC<sub>50</sub>(log-mean), was preserved in the replicate dataset.
